# gtfsort: a tool to efficiently sort GTF files

**DOI:** 10.1101/2023.10.21.563454

**Authors:** Alejandro Gonzales-Irribarren, Anne Fu

## Abstract

The General Transfer Format (GTF) is a widely used format for gene annotation data, integral to various downstream analyses. Efficient management and sorting of GTF data are relevant, as unsorted data can intere with computational efficiency and interpretability. Sorted GTF data, on the other hand, enable more intuitive navigation of genomic structures and can enhance the performance of some bioinformatics tools. This paper presents gtfsort, a tool written in Rust, which efficiently sorts GTF files, outperforming existing software. By reducing computation times and providing structured outputs, gtfsort enhances the interpretation, relationships, and readability of gene annotation data.

## Background

The General Transfer Format (GTF) is one of the primary and widely used formats when working with gene annotation data, employed to store information about gene structure [1]. In GTF files, details regarding gene features, exon boundaries, and various attributes associated with genomic elements are encapsulated, although not necessarily in an organized manner [2]. As the foundation for numerous downstream analyses, the effective management and sorting of GTF data have emerged as a challenge. Unsorted GTF data can significantly impact computation times and hinder the efficiency of these downstream tools, especially when conducting tasks such as locating specific gene features, extracting relevant information, or comparing data across multiple genes and transcripts [3, 4].

On the other hand, sorted GTF data can substantially improve the performance of bioinformatics tools and software, simplifying the retrieval of specific genomic elements and reducing resources required for data access [5, 6]. Furthermore, the organization of GTF data into a coherent and systematic structure offers users a more intuitive and natural perspective on gene structures. Therefore, showing these information as a logical sequence, allowing researchers to traverse through genomic elements with ease [5]. This structured view enhances the interpretability of gene annotation data, making it simpler to discern relationships between features, exons, and transcripts [2]. While different sorting approaches could be applied over a GTF file, sorting only by the start position, for example; the most complete procedure will need to account for chromosome position, start coordinates and features.

This paper presents gtfsort, a sorting tool that utilizes a lexicographically-based index ordering algorithm and holds the potential to address this challenge. By offering an efficient means of sorting GTF data, gtfsort not only outperforms similar tools such as GFF3sort [5] or AGAT [7] but also provides a more natural, ordered, and user-friendly perspective on GTF structure.

### Implementation

This tool is written in Rust and presented as a standalone command-line tool and a library. The user only needs to specific input GTF and output GTF files. Upon reading the input file, gtfsort organizes the data into three distinct layers: an outer layer for the highest-level hierarchy, an inner layer for lower-level hierarchies, and a transcript-mapper layer responsible for managing isoforms and their associated features for a given gene. Each line in the GTF file is parsed and grouped according to its feature, aligning with the specific layer-dependent data flow, including genes, transcripts, and lower-level hierarchies. The parsing structure of the grandparent node implements a quick access to 3 key variables for the sorting step: chromosome, start position and gene id, that are pushed to a vector. At this point, where each line in the GTF file has been matched with a feature and located at their specific layer, gtfsort just orders the outer vector by chromosome and start position (Fig.1A).

**Figure 1:**
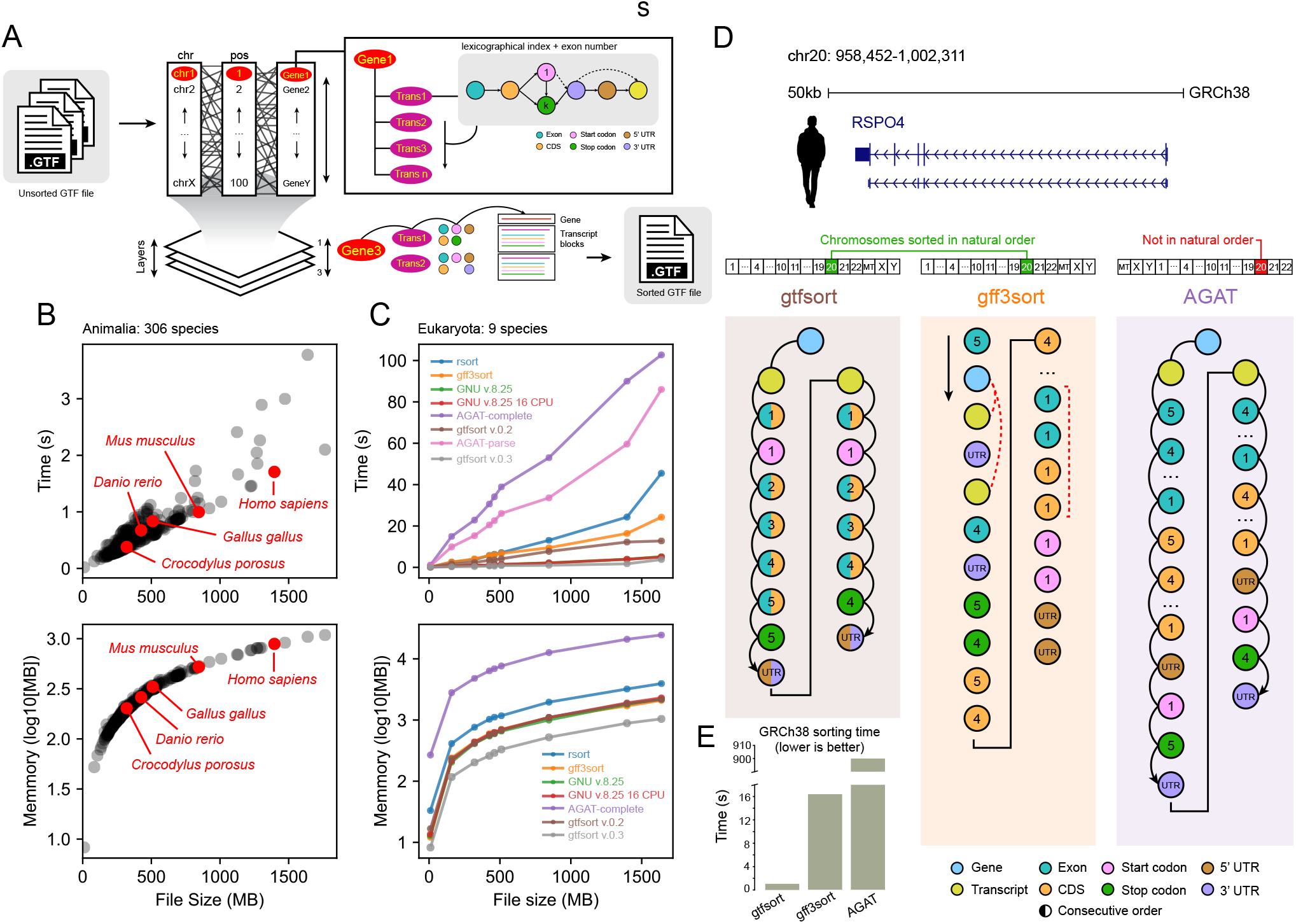
gtfsort outperforms existing sorting tools. **A**. Implementing a lexicographically-based index algorithm, gtfsort utilizes multiple layers to efficiently write transcript blocks. **B**. The Ensembl Animalia dataset (306 species) sorted with gtfsort, time/memmory usage are highlighted for reference species. The biggest annotation file (*Cyprinus carpio carpio* 1.9 Gb) takes around 3 seconds and 1 Gbs of RAM. **C**. Benchmarking 7 different sorting tools using 9 Eukaryota species. gtfsort v.0.3 is the lowest time and and less memmory consumer tool. AGAT-complete and AGAT-parse values are presented as their decimal part for visualization purposes. **D**. RSPO4 sorting output comparison between GFF3sort, AGAT and gtfsort. gtfsort is the only tool with intuitive outputs at chromosome and feature ordering levels. **E**. Sorting time comparison between GFF3sort, AGAT and gtfsort. gtfsort v.0.3 is the fastest tool, with 1.71 seconds.

The transcript-mapper layer plays a key role after that, supported by a helper hash storing gene:transcript pairs, is the binding between a gene an a transcript block (transcript + childs). In short, for a given gene *j* with *n* transcripts, where *n* ∈ *n*_1_, *n*_2_, …, *n*_*k*_ and *k* is the upper bound range of transcripts for *j*, each *n* is the key for all low-level features stored in the inner-layer, a lexicographically ordered hash. Here, exon numbers and specific feature matches work in tandem, creating a directed cycle. This allows an in-site sort for all attributes that are accessed by the transcript key, drastically reducing the complexity. Finally, gtfsort just loops through the first and second layers to retrieve all blocks and directly write them to the output file (Fig.1A).

To assess the efficiency and results of gtfsort, two main benchmarks were conducted. Initially, gtfsort (v.0.2 and v.0.3) was applied to the entire Ensembl Animalia GTF3 dataset (110th release; 306 species) [8]. Subsequently, a comparative analysis of the gtfsort utility in relation to several existing software tools: GNU v.8.25 (both in single and multi-core configurations) [9], AGAT (utilizing the –gff flag for both complete and partial parsing phases) [7], GFF3sort (with specific options, including –precise and –chr_order natural) [5], and rsort (an unpublished multicore Rust implementation with nested data structures) was conducted. This comprehensive evaluation encompassed a diverse array of biological domains, spanning bacteria, fungi, insects, mammals, and more. To ensure a robust assessment, nine common species were selected: *Homo sapiens, Mus musculus, Canis lupus familiaris, Gallus gallus, Danio rerio, Salmo salar, Crocodylus porosus, Drosophila melanogaster* and *Saccharomyces cerevisiae*. All tests are presented as the mean of 10 runs and were performed on a 5700X AMD CPU, 128 Gbs RAM, running Fedora 37.

From the suite of tools employed in the preceding step, only three assert to incorporate a feature sorting step: GFF3sort, AGAT, and gtfsort. Both, GFF3sort and AGAT, employ a topological algorithm to build directed acyclic graphs of parent-child relations that guides the ordering sense. To determine the real sorting performance of these three tools, that are announced as specific software to sort gene annotation data, two variables were chosen: GRCh38 sorting time and the actual line-by-line sorting output. For the latter, a random gene with at least two isoforms was selected.

## Results and Discussion

In the first benchmark, by sorting a 1.9 Gbs GTF file (*Cyprinus carpio carpio*) in 2.1 seconds and less than 1 Gb of RAM with high accuracy, gtfsort demonstrated both of their main attributes: speed and efficiency. The relation between the file size and time is linear, consistent with the memory usage by the process. Since *Cyprinus carpio carpio* is the species with the heaviest annotation file, its score is taken as the upper bound of gtfsort capabilities. Other species such as *Mus musculus* or *Gallus gallus*, of greater importance in medicine and biology, can be handled in less than 1 second and using a minimum of 0.5 Gbs of RAM by gtfsort (Fig.1B).

The second benchmark compared 7 tools, between unique tools or tools in different modes. Here, gtfsort demonstrated remarkable efficiency, showcasing the shortest computation time along with the GNU software (in both single and multi-core modes). It is worth noting, however, that GNU software fails to consistently maintain a stable chromosome/position/feature order and encounters difficulties when sorting commented lines (e.g., lines commencing with ‘#’ at the beginning of the file). The remaining tools exhibited substantially longer processing times, with some employing parallel processing approaches (e.g. rsort, which utilized 16 cores). Furthermore, is also notable that the less memory allocation required for sorting files remained conservative in three of the tools evaluated: GNU (both single and multi-core), GFF3sort, and gtfsort. The memory utilization for the largest file did not exceed 2.3 Gbs, even when handling substantial datasets (up to 1.6 Gbs in size) (Fig.1C).

Among the tools compared in the final test, gtfsort emerged as the fastest with a processing time of 1.71 seconds, followed by GFF3sort at 16.40 seconds, and AGAT, which required approximately 900 seconds to complete the sorting operation (Fig.1E). The notorious difference with the extensive computation time of AGAT is due to the fact that does not only sort a GTF file but inspects some controversial lines and fixes/adds corrected/missing lines. Furthermore, when inspecting the actual sorting output at a chromosomal ordering level, this study found that only gtfsort and GFF3sort present an intuitive ordering (starting with chr1 and ending with chrX). AGAT fails here, locating chrMT and both sex chromosomes first. At a feature level, nevertheless, GFF3sort failed to present an ordered structure of features. This is quickly perceived since the first line is occupied by an “exon” feature instead the grandparent node: the “gene” feature. In contrast, AGAT and gtfsort, do exhibit an intuitive structure order, initiating with a “gene” feature followed by the first “transcript” feature and a low-hierarchy attributes next. AGAT presents 2 blocks per transcript, all CDS after all exons with start/stop codons and UTRs at the end. gtfsort, on the other hand, adopted a distinct approach, presenting pairs or triplets of features in conjunction with their respective exon numbers, sorted in descending order, even for sequences on negative strands. UTRs were consistently positioned at the conclusion of the sequence, enabling a natural and rapid comprehension of the information associated with a given exon (exon/CDS/start/stop) (Fig.1D).

## Conclusions

This paper introduces gtfsort, a sorting tool that implements a lexicographically-based index algorithm outperforming similar existing software, making it a valuable addition to the bioinformatics toolkit. By reducing computation times and providing an accurate and structured output, this tool will help to make gene annotation data easier to interpret, relate and read by the user.

## Availability and requirements

Project name: gtfsort

Project home page: github.com/alejandrogzi/gtfsort

Operating system(s): Linux, MacOS, Windows

Programming language: Rust

Other requirements: No License: MIT

