## Supplemental Material for "gtfsort: a tool to efficiently sort GTF files"

### Sofware

GNU (v.8.25) [1]: `sort -k1,1 -k4,4n -k5,5n file.gtf > out.gtf`
GNU 16 (v. 8.25) [1]: `sort -k1,1 -k4,4n -k5,5n -parallel 16 file.gtf > out.gtf`
rsort (v. 0.1.0) (this paper): `rsort file.gtf out.gtf -t 16`
AGAT (v. 1.0.0) [2]: `agat_convert_sp_gxf2gxf.pl -gff file.gtf`
GFF3sort (v. 1.0.0) [3]: `gff3sort.pl -precise -chr_order natural file.gtf > out.gtf`
gtfsort (v. 0.3.0) (this paper): `gtfsort file.gtf out.gtf`

### Benchmark Data

Table S1: Comparison of the running time (s) of different GTF sorting tools

| *Species* | Assembly[1] | rsort | gff3sort | GNU | GNU 16 | AGATc[2] | gtfsort v0.2 | AGATp[3] | gtfsort v0.3 |
|---|---|---|---|---|---|---|---|---|---|
| *H. sapiens* | GRCh38 | 24.31 | 16.40 | 3.92 | 3.89 | 900.00 | 12.02 | 596.00 | 1.71 |
| *M. musculus* | GRCm39 | 13.08 | 9.48 | 2.17 | 2.11 | 530.08 | 7.71 | 336.00 | 0.99 |
| *C. familiaris* | ROS_Cfam_1.0 | 6.34 | 6.10 | 1.35 | 1.26 | 341.33 | 3.62 | 225.62 | 0.59 |
| *G. gallus* | bGalGal1 | 7.13 | 6.74 | 1.41 | 1.42 | 388.98 | 4.14 | 260.88 | 0.84 |
| *S. cerevisiae* | R64-1-1 | 0.20 | 0.22 | 0.11 | 0.11 | 11.04 | 0.16 | 7.30 | 0.02 |
| *D. rerio* | GRCz11 | 5.78 | 5.72 | 1.03 | 1.05 | 306.18 | 3.43 | 205.34 | 0.67 |
| *D. melanogaster* | BDGP6.46 | 2.50 | 2.55 | 0.61 | 0.54 | 149.63 | 1.58 | 98.90 | 0.28 |
| *C. porosus* | CroPor_comp1 | 3.67 | 4.12 | 1.10 | 1.06 | 228.45 | 2.36 | 153.21 | 0.38 |
| *S. salar* | Ssal_v3 | 45.50 | 24.26 | 5.15 | 5.02 | 1027.24 | 12.76 | 860.23 | 3.78 |

Table S2: Comparison of the memmory usage (s) of different GTF sorting tools

| *Species* | Assembly[1] | rsort | gff3sort | GNU | GNU 16 | AGAT | gtfsort v0.2 | gtfsort v0.3 |
|---|---|---|---|---|---|---|---|---|
| *H. sapiens* | GRCh38 | 3216.74 | 1700.00 | 1800.00 | 1900.00 | 20 900.00 | 1850.39 | 886.45 |
| *M. musculus* | GRCm39 | 1959.18 | 1100.00 | 1000.00 | 1100.00 | 12 658.48 | 1102.69 | 521.92 |
| *C. familiaris* | ROS_Cfam_1.0 | 1112.58 | 625.60 | 594.48 | 627.51 | 6902.61 | 604.92 | 288.96 |
| *G. gallus* | bGalGal1 | 1176.13 | 705.00 | 657.85 | 694.40 | 7638.35 | 680.31 | 330.92 |
| *S. cerevisiae* | R64-1-1 | 33.15 | 12.15 | 12.87 | 13.58 | 268.20 | 16.75 | 8.25 |
| *D. rerio* | GRCz11 | 1028.83 | 600.50 | 548.76 | 579.25 | 6371.77 | 563.73 | 260.69 |
| *D. melanogaster* | BDGP6.46 | 412.31 | 241.00 | 205.18 | 216.58 | 2800.00 | 232.21 | 117.11 |
| *C. porosus* | CroPor_comp1 | 763.42 | 439.20 | 413.28 | 436.24 | 4798.64 | 413.94 | 202.34 |
| *S. salar* | Ssal_v3 | 3939.62 | 2100.00 | 2200.00 | 2300.00 | 24 530.69 | 2167.52 | 1043.63 |

[1] All annotation files were downloaded from Ensembl [4].
[2] AGATc: AGAT-complete
[3] AGATp: AGAT-parse

### Sorting Output

End positions, frames and miscellaneous attributes have been removed for clarity.

GFF3sort (v. 1.0.0) [3]:

```
20 ensembl_havana exon 958452  gene_id ‘ENSG00000101282’; transcript_id ‘ENST00000217260’; exon_number ‘5’;
20 ensembl_havana gene 958452  gene_id ‘ENSG00000101282’;
20 ensembl_havana transcript 958452  gene_id ‘ENSG00000101282’; transcript_id ‘ENST00000217260’;
20 ensembl_havana three_prime_utr 958452  gene_id ‘ENSG00000101282’; transcript_id ‘ENST00000217260’;
20 ensembl_havana transcript 960254  gene_id ‘ENSG00000101282’; transcript_id ‘ENST00000400634’;
20 ensembl_havana exon 960254  gene_id ‘ENSG00000101282’; transcript_id ‘ENST00000400634’; exon_number ‘4’;
20 ensembl_havana three_prime_utr 960254  gene_id ‘ENSG00000101282’; transcript_id ‘ENST00000400634’;
20 ensembl_havana stop_codon 960357  gene_id ‘ENSG00000101282’; transcript_id ‘ENST00000217260’; exon_number ‘5’;
20 ensembl_havana stop_codon 960357  gene_id ‘ENSG00000101282’; transcript_id ‘ENST00000400634’; exon_number ‘4’;
20 ensembl_havana CDS 960360  gene_id ‘ENSG00000101282’; transcript_id ‘ENST00000217260’; exon_number ‘5’;
20 ensembl_havana CDS 960360  gene_id ‘ENSG00000101282’; transcript_id ‘ENST00000400634’; exon_number ‘4’;
20 ensembl_havana exon 963935  gene_id ‘ENSG00000101282’; transcript_id ‘ENST00000217260’; exon_number ‘4’;
20 ensembl_havana CDS 963935  gene_id ‘ENSG00000101282’; transcript_id ‘ENST00000217260’; exon_number ‘4’;
20 ensembl_havana exon 967174  gene_id ‘ENSG00000101282’; transcript_id ‘ENST00000400634’; exon_number ‘3’;
20 ensembl_havana exon 967174  gene_id ‘ENSG00000101282’; transcript_id ‘ENST00000217260’; exon_number ‘3’;
20 ensembl_havana CDS 967174  gene_id ‘ENSG00000101282’; transcript_id ‘ENST00000400634’; exon_number ‘3’;
20 ensembl_havana CDS 967174  gene_id ‘ENSG00000101282’; transcript_id ‘ENST00000217260’; exon_number ‘3’;
20 ensembl_havana exon 967950  gene_id ‘ENSG00000101282’; transcript_id ‘ENST00000400634’; exon_number ‘2’;
20 ensembl_havana CDS 967950  gene_id ‘ENSG00000101282’; transcript_id ‘ENST00000400634’; exon_number ‘2’;
20 ensembl_havana CDS 967950  gene_id ‘ENSG00000101282’; transcript_id ‘ENST00000217260’; exon_number ‘2’;
20 ensembl_havana exon 967950  gene_id ‘ENSG00000101282’; transcript_id ‘ENST00000217260’; exon_number ‘2’;
20 ensembl_havana exon 1002086  gene_id ‘ENSG00000101282’; transcript_id ‘ENST00000400634’; exon_number ‘1’;
20 ensembl_havana CDS 1002086  gene_id ‘ENSG00000101282’; transcript_id ‘ENST00000400634’; exon_number ‘1’;
20 ensembl_havana exon 1002086  gene_id ‘ENSG00000101282’; transcript_id ‘ENST00000217260’; exon_number ‘1’;
20 ensembl_havana CDS 1002086  gene_id ‘ENSG00000101282’; transcript_id ‘ENST00000217260’; exon_number ‘1’;
20 ensembl_havana start_codon 1002162  gene_id ‘ENSG00000101282’; transcript_id ‘ENST00000400634’; exon_number ‘1’;
20 ensembl_havana start_codon 1002162  gene_id ‘ENSG00000101282’; transcript_id ‘ENST00000217260’; exon_number ‘1’;
20 ensembl_havana five_prime_utr 1002165  gene_id ‘ENSG00000101282’; transcript_id ‘ENST00000400634’;
20 ensembl_havana five_prime_utr 1002165  gene_id ‘ENSG00000101282’; transcript_id ‘ENST00000217260’;
```

AGAT (v. 1.0.0) [2]:

```
20 ensembl_havana gene 958452 ID=ENSG00000101282;gene_id=ENSG00000101282;
20 ensembl_havana transcript 958452 ID=ENST00000217260;Parent=ENSG00000101282;
20 ensembl_havana exon 958452 ID=ENSE00000990907;Parent=ENST00000217260;exon_number=5;
20 ensembl_havana exon 963935 ID=ENSE00000858560;Parent=ENST00000217260;exon_number=4;
20 ensembl_havana exon 967174 ID=ENSE00000655139;Parent=ENST00000217260;exon_number=3;
20 ensembl_havana exon 967950 ID=ENSE00000858561;Parent=ENST00000217260;exon_number=2;
20 ensembl_havana exon 1002086 ID=ENSE00000858562;Parent=ENST00000217260;exon_number=1;
20 ensembl_havana CDS 960357 ID=cds-785868;Parent=ENST00000217260;exon_number=5;
20 ensembl_havana CDS 963935 ID=cds-785867;Parent=ENST00000217260;exon_number=4;
20 ensembl_havana CDS 967174 ID=cds-785866;Parent=ENST00000217260;exon_number=3;
20 ensembl_havana CDS 967950 ID=cds-785865;Parent=ENST00000217260;exon_number=2;
20 ensembl_havana CDS 1002086 ID=cds-785864;Parent=ENST00000217260;exon_number=1;
20 ensembl_havana five_prime_utr ID=five_prime_utr-151948;Parent=ENST00000217260;
20 ensembl_havana start_codon 1002162 ID=start_codon-85688;Parent=ENST00000217260;exon_number=1;
20 ensembl_havana stop_codon 960357 ID=stop_codon-80449;Parent=ENST00000217260;exon_number=5;
20 ensembl_havana three_prime_utr 958452 ID=three_prime_utr-185450;Parent=ENST00000217260;
20 ensembl_havana transcript 960254 ID=ENST00000400634;Parent=ENSG00000101282;
20 ensembl_havana exon 960254 ID=ENSE00001855250;Parent=ENST00000400634;exon_number=4;
20 ensembl_havana exon 967174 ID=nbis-exon-761509;Parent=ENST00000400634;exon_number=3;
20 ensembl_havana exon 967950 ID=nbis-exon-761508;Parent=ENST00000400634;exon_number=2;
20 ensembl_havana exon 1002086 ID=ENSE00001842772;Parent=ENST00000400634;exon_number=1;
20 ensembl_havana CDS 960357 ID=cds-785872;Parent=ENST00000400634;exon_number=4;
20 ensembl_havana CDS 967174 ID=cds-785871;Parent=ENST00000400634;exon_number=3;
20 ensembl_havana CDS 967950 ID=cds-785870;Parent=ENST00000400634;exon_number=2;
20 ensembl_havana CDS 1002086 ID=cds-785869;Parent=ENST00000400634;exon_number=1;
20 ensembl_havana five_prime_utr 1002165 ID=five_prime_utr-151949;Parent=ENST00000400634;
20 ensembl_havana start_codon 1002162 ID=start_codon-85689;Parent=ENST00000400634;exon_number=1;
20 ensembl_havana transcript 960466 ID=ENST00000474461;Parent=ENSG00000101282;
20 ensembl_havana exon 1002086 ID=ENSE00001884975;Parent=ENST00000474461;exon_number=1;
```

```
20 ensembl_havana CDS 1002086 ID=cds-785874;Parent=ENST00000474461;exon_number=1;
```

gtfsort (v. 0.3.0) (this paper):

```
20 ensembl_havana gene 958452  gene_id "ENSG00000101282";
20 ensembl_havana transcript 958452  gene_id "ENSG00000101282"; transcript_id "ENST00000217260";
20 ensembl_havana exon 1002086  gene_id "ENSG00000101282"; transcript_id "ENST00000217260"; exon_number "1";
20 ensembl_havana CDS 1002086  gene_id "ENSG00000101282"; transcript_id "ENST00000217260"; exon_number "1";
20 ensembl_havana start_codon 1002162  gene_id "ENSG00000101282"; transcript_id "ENST00000217260"; exon_number "1";
20 ensembl_havana exon 967950  gene_id "ENSG00000101282"; transcript_id "ENST00000217260"; exon_number "2";
20 ensembl_havana CDS 967950  gene_id "ENSG00000101282"; transcript_id "ENST00000217260"; exon_number "2";
20 ensembl_havana exon 967174  gene_id "ENSG00000101282"; transcript_id "ENST00000217260"; exon_number "3";
20 ensembl_havana CDS 967174  gene_id "ENSG00000101282"; transcript_id "ENST00000217260"; exon_number "3";
20 ensembl_havana exon 963935  gene_id "ENSG00000101282"; transcript_id "ENST00000217260"; exon_number "4";
20 ensembl_havana CDS 963935  gene_id "ENSG00000101282"; transcript_id "ENST00000217260"; exon_number "4";
20 ensembl_havana exon 958452  gene_id "ENSG00000101282"; transcript_id "ENST00000217260"; exon_number "5";
20 ensembl_havana CDS 960360  gene_id "ENSG00000101282"; transcript_id "ENST00000217260"; exon_number "5";
20 ensembl_havana stop_codon 960357  gene_id "ENSG00000101282"; transcript_id "ENST00000217260"; exon_number "5";
20 ensembl_havana transcript 960254  gene_id "ENSG00000101282"; transcript_id "ENST00000400634";
20 ensembl_havana exon 1002086  gene_id "ENSG00000101282"; transcript_id "ENST00000400634"; exon_number "1";
20 ensembl_havana CDS 1002086  gene_id "ENSG00000101282"; transcript_id "ENST00000400634"; exon_number "1";
20 ensembl_havana start_codon 1002162  gene_id "ENSG00000101282"; transcript_id "ENST00000400634"; exon_number "1";
20 ensembl_havana exon 967950  gene_id "ENSG00000101282"; transcript_id "ENST00000400634"; exon_number "2";
20 ensembl_havana CDS 967950  gene_id "ENSG00000101282"; transcript_id "ENST00000400634"; exon_number "2";
20 ensembl_havana exon 967174  gene_id "ENSG00000101282"; transcript_id "ENST00000400634"; exon_number "3";
20 ensembl_havana CDS 967174  gene_id "ENSG00000101282"; transcript_id "ENST00000400634"; exon_number "3";
20 ensembl_havana exon 960254  gene_id "ENSG00000101282"; transcript_id "ENST00000400634"; exon_number "4";
20 ensembl_havana CDS 960360  gene_id "ENSG00000101282"; transcript_id "ENST00000400634"; exon_number "4";
20 ensembl_havana stop_codon 960357  gene_id "ENSG00000101282"; transcript_id "ENST00000400634"; exon_number "4";
20 ensembl_havana five_prime_utr 1002165  gene_id "ENSG00000101282"; transcript_id "ENST00000400634";
20 ensembl_havana three_prime_utr 960254  gene_id "ENSG00000101282"; transcript_id "ENST00000400634";
```

### References


[1] Haertel Mike and Eggert Paul. *GNU Core Utilities*. Free Software Foundation, 2016.

[2] Jacques Dainat, Darío Hereñú, KD. Murray, Ed Davis, Kathryn Crouch, Lucile Sol, Nuno Agostinho, and Zachary Zollman. AGAT: Another Gff Analysis Toolkit to handle annotations in any GTF/GFF format. *Zenodo*, 2023.

[3] Tao Zhu, Chengzhen Liang, Zhigang Meng, Sandui Guo, and Rui Zhang. GFF3sort: a novel tool to sort GFF3 files for tabix indexing. *BMC bioinformatics*, 18:1–4, 2017.

[4] Fiona Cunningham, James E Allen, Jamie Allen, Jorge Alvarez-Jarreta, M Ridwan Amode, Irina M Armean, Olanrewaju Austine-Orimoloye, Andrey G Azov, If Barnes, Ruth Bennett, et al. Ensembl 2022. *Nucleic acids research*, 50(D1):D988–D995, 2022.
